## Supplemental Table 1 for "The small RNA component of Arabidopsis phloem sap and its response to iron deficiency"

Table 1. List of over- and under-represented tRNA genes in phloem exudates

| tRNA | Fold Change | p-value |
| --- | --- | --- |
| HisGTG.4 | 126.8087 | 4.60E-111 |
| HisGTG.9 | 124.521 | 4.66E-105 |
| GluTTC.9 | 123.596 | 1.33E-78 |
| HisGTG.7 | 117.4119 | 1.24E-103 |
| HisGTG.1 | 112.9794 | 6.10E-103 |
| HisGTG.8 | 109.567 | 7.11E-106 |
| HisGTG.2 | 104.928 | 5.78E-95 |
| HisGTG.10 | 85.77149 | 2.31E-65 |
| HisGTG.6 | 82.69842 | 6.28E-141 |
| AspGTC.8 | 81.33028 | 2.37E-46 |
| IleAAT.3 | 80.82767 | 1.83E-37 |
| HisGTG.5 | 74.91785 | 4.21E-75 |
| AlaCGC.1 | 73.6072 | 1.57E-73 |
| HisGTG.3 | 73.33672 | 7.18E-75 |
| GlyTCC.10 | 71.21723 | 2.42E-77 |
| GlyTCC.1 | 71.12381 | 4.08E-78 |
| GlyTCC.8 | 70.86892 | 4.79E-78 |
| GlyTCC.5 | 70.76431 | 3.01E-78 |
| GlyTCC.3 | 70.30388 | 5.03E-78 |
| GlyTCC.7 | 70.18915 | 1.01E-77 |
| GlyTCC.11 | 70.10081 | 9.69E-78 |
| GlyTCC.9 | 69.90008 | 1.94E-77 |
| GlyTCC.6 | 69.6128 | 3.28E-77 |
| GlyTCC.2 | 69.30529 | 1.87E-77 |
| AspGTC.26 | 66.32611 | 3.49E-42 |
| GlyTCC.4 | 66.08256 | 5.44E-76 |
| GlyTCC.12 | 65.05141 | 1.34E-75 |
| AspGTC.15 | 64.02325 | 3.55E-41 |
| AspGTC.16 | 61.4105 | 3.11E-39 |
| AlaCGC.5 | 60.25187 | 1.62E-78 |
| AspGTC.12 | 60.22719 | 6.31E-41 |
| AspGTC.25 | 57.89545 | 9.70E-42 |
| LeuTAG.8 | 56.87934 | 1.82E-86 |
| AspGTC.2 | 56.11129 | 2.80E-41 |
| AspGTC.14 | 55.88649 | 2.58E-38 |
| AspGTC.1 | 54.82048 | 3.46E-42 |
| AspGTC.7 | 52.85105 | 3.64E-43 |
| AspGTC.23 | 52.33664 | 2.32E-36 |
| AspGTC.4 | 52.21638 | 3.53E-39 |
| AlaCGC.6 | 51.43856 | 2.99E-73 |
| AlaCGC.2 | 49.6925 | 9.56E-73 |
| AspGTC.20 | 49.18953 | 3.87E-40 |
| AspGTC.10 | 49.12048 | 7.51E-39 |
| AspGTC.9 | 48.89438 | 6.52E-39 |
| ThrAGT.6 | 48.80215 | 3.73E-60 |
| AspGTC.17 | 48.12788 | 8.27E-37 |
| AlaCGC.4 | 48.06095 | 3.29E-59 |
| AspGTC.19 | 45.22254 | 1.01E-34 |
| UndUND.1 | 45.12357 | 3.00E-71 |
| CysGCA.1 | 45.00804 | 4.76E-30 |
| AspGTC.3 | 42.67274 | 4.22E-38 |
| AspGTC.11 | 42.00936 | 4.55E-39 |
| AspGTC.21 | 40.25189 | 3.15E-35 |
| AspGTC.24 | 38.53381 | 1.93E-36 |
| AspGTC.13 | 37.69307 | 3.29E-83 |
| GluTTC.8 | 35.93363 | 3.80E-32 |
| GluTTC.3 | 35.88659 | 2.99E-32 |
| GluTTC.7 | 30.84394 | 3.82E-32 |
| AspGTC.6 | 29.69997 | 4.02E-29 |
| AspGTC.18 | 28.8241 | 3.80E-29 |
| LysTTT.3 | 24.00596 | 2.31E-09 |
| ArgCCG.4 | 22.87971 | 4.31E-17 |
| SerGCT.5 | 20.73254 | 2.65E-85 |
| MetCAT.8 | 17.999 | 7.89E-25 |
| ValCAC.2 | 17.52892 | 1.20E-64 |
| SerGCT.2 | 17.20511 | 2.62E-62 |
| SerTGA.8 | 13.44203 | 1.07E-23 |
| ArgCCG.5 | 13.29096 | 1.40E-12 |
| ArgCCT.5 | 12.10052 | 1.77E-11 |
| ArgCCG.3 | 11.9533 | 2.74E-11 |
| GluTTC.2 | 11.73497 | 4.91E-14 |
| GluTTC.13 | 11.61349 | 3.87E-14 |
| GluTTC.12 | 11.58996 | 4.96E-14 |
| GluTTC.10 | 11.55837 | 6.14E-14 |
| GluTTC.14 | 11.55418 | 6.31E-14 |
| LysCTT.14 | 11.43656 | 0.000152631 |
| AsnGTT.6 | 10.7327 | 2.21E-28 |
| SerGGA.1 | 10.55566 | 5.52E-31 |
| GluTTC.4 | 10.50725 | 3.70E-13 |
| SerTGA.7 | 9.736465 | 1.87E-17 |
| ThrAGT.3 | 9.204248 | 3.83E-12 |
| CysGCA.13 | 8.363118 | 1.10E-16 |
| ProTGG.37 | 8.182341 | 1.12E-12 |
| MetCAT.13 | 7.960326 | 4.43E-12 |
| TyrGTA.59 | 7.55414 | 1.18E-13 |
| GluCTC.9 | 7.253044 | 4.45E-13 |
| TyrGTA.68 | 7.025169 | 1.78E-13 |
| GluCTC.8 | 7.015465 | 1.58E-12 |
| ThrAGT.4 | 6.929147 | 1.16E-11 |
| ThrAGT.1 | 6.883453 | 3.30E-09 |
| GlyGCC.8 | 6.623836 | 4.08E-10 |
| ThrAGT.5 | 6.548199 | 9.39E-06 |
| GlyGCC.3 | 6.323454 | 5.94E-07 |
| ArgTCG.5 | 6.008941 | 2.59E-14 |
| AlaAGC.1 | 5.799297 | 3.25E-17 |
| ArgTCG.4 | 5.703401 | 4.70E-12 |
| AlaAGC.8 | 5.628906 | 7.05E-15 |
| ArgTCG.3 | 5.575474 | 4.95E-12 |
| TyrGTA.57 | 5.513751 | 9.30E-06 |
| AlaAGC.15 | 5.409308 | 3.77E-14 |
| AlaAGC.12 | 5.315495 | 1.65E-14 |
| AlaAGC.13 | 5.314556 | 3.14E-12 |
| AlaAGC.7 | 5.298362 | 3.46E-14 |
| AlaAGC.10 | 5.275042 | 2.53E-15 |
| GlyACC.1 | 5.271454 | 2.36E-05 |
| AlaAGC.14 | 5.212589 | 6.54E-14 |
| CysGCA.4 | 5.178631 | 6.06E-07 |
| AlaAGC.11 | 5.159728 | 6.25E-13 |
| AlaAGC.5 | 5.157423 | 6.79E-14 |
| AlaAGC.6 | 5.151627 | 2.66E-14 |
| AlaAGC.2 | 5.140755 | 1.29E-14 |
| IleAAT.2 | 5.116291 | 0.026570781 |
| AlaAGC.3 | 5.108752 | 1.13E-11 |
| GlyGCC.6 | 5.083897 | 3.14E-07 |
| AlaAGC.9 | 5.013226 | 1.59E-12 |
| MetCAT.14 | 4.922102 | 1.59E-10 |
| AlaAGC.16 | 4.868087 | 1.38E-11 |
| GlnTTG.5 | 4.864521 | 0.000776722 |
| TrpCCA.6 | 4.738755 | 1.25E-15 |
| ArgTCG.6 | 4.595499 | 1.30E-07 |
| ThrAGT.7 | 4.43894 | 0.003504094 |
| SerTGA.1 | 4.426341 | 9.54E-07 |
| MetCAT.10 | 4.357604 | 0.020583544 |
| GluTTC.6 | 4.259999 | 3.17E-07 |
| GlyGCC.9 | 3.873466 | 0.003717233 |
| GlyGCC.14 | 3.807823 | 0.007257673 |
| AlaTGC.5 | 3.79967 | 3.22E-06 |
| AlaTGC.1 | 3.760419 | 3.16E-05 |
| AlaTGC.4 | 3.673484 | 4.47E-06 |
| GlyGCC.7 | 3.636475 | 7.15E-05 |
| AlaTGC.2 | 3.635021 | 7.86E-06 |
| ArgCCT.6 | 3.612126 | 0.006515685 |
| AlaTGC.6 | 3.595868 | 3.28E-06 |
| AlaAGC.4 | 3.572196 | 4.55E-05 |
| AlaTGC.9 | 3.566412 | 6.92E-05 |
| AlaTGC.7 | 3.473683 | 8.08E-05 |
| CysGCA.3 | 3.442598 | 0.019603472 |
| GlyGCC.1 | 3.398198 | 0.001290906 |
| ArgCCT.8 | 3.339192 | 0.001556018 |
| PheGAA.5 | 3.320609 | 0.000376184 |
| IleAAT.8 | 3.272842 | 0.013619012 |
| ThrAGT.9 | 3.2155 | 0.012934559 |
| GluCTC.11 | 3.148056 | 0.0019239 |
| ArgTCG.2 | 3.144542 | 0.001380552 |
| GluCTC.10 | 3.074907 | 0.002922135 |
| GluCTC.2 | 3.068699 | 0.003064918 |
| TyrGTA.66 | 3.042184 | 0.009373644 |
| GluCTC.12 | 3.01227 | 0.003827711 |
| GluCTC.3 | 3.008133 | 0.004357549 |
| AspGTC.22 | 3.005197 | 0.035712591 |
| GluCTC.13 | 2.957438 | 0.004921427 |
| GluCTC.6 | 2.955582 | 0.009086004 |
| GluCTC.4 | 2.953555 | 0.006453426 |
| GluCTC.1 | 2.953498 | 0.008209847 |
| GluCTC.7 | 2.943364 | 0.005320641 |
| IleAAT.10 | 2.935611 | 0.037927982 |
| AlaTGC.8 | 2.856308 | 0.003405372 |
| AlaCGC.3 | 2.784761 | 0.001974717 |
| ValAAC.15 | -2.7076 | 0.015428728 |
| ProAGG.3 | -2.89065 | 0.049895835 |
| GlnCTG.3 | -2.91054 | 0.00962104 |
| TrpCCA.3 | -2.96782 | 0.00193786 |
| GlnCTG.4 | -2.96849 | 0.0026138 |
| GlyGCC.10 | -2.99039 | 0.024889627 |
| ProTGG.11 | -3.00719 | 0.018344298 |
| LysTTT.14 | -3.08846 | 0.028588268 |
| ThrCGT.4 | -3.10403 | 0.014260698 |
| TyrGTA.74 | -3.19309 | 0.015759774 |
| ProAGG.12 | -3.22373 | 0.017083218 |
| TyrGTA.70 | -3.36232 | 0.007472825 |
| SerTGA.4 | -3.36705 | 0.005140779 |
| GlyGCC.24 | -3.3932 | 0.006385935 |
| SerTGA.3 | -3.45182 | 0.007452487 |
| GlnCTG.6 | -3.46793 | 0.000603946 |
| TyrGTA.67 | -3.65062 | 0.010276463 |
| LeuTAG.4 | -3.68527 | 0.000152265 |
| GlnCTG.8 | -3.71425 | 3.64E-05 |
| TyrGTA.76 | -3.78218 | 0.01822488 |
| LeuTAG.9 | -3.79657 | 0.002136997 |
| ThrTGT.5 | -3.80729 | 2.28E-06 |
| GlyGCC.18 | -3.81756 | 0.000102931 |
| CysGCA.10 | -3.87685 | 0.004720772 |
| AsnGTT.9 | -4.01915 | 0.000850354 |
| TyrGTA.73 | -4.14825 | 3.49E-06 |
| LeuTAG.1 | -4.20134 | 0.000157817 |
| LeuTAG.6 | -4.2946 | 0.001487155 |
| TyrGTA.61 | -4.5122 | 0.000103571 |
| LeuTAG.2 | -4.52191 | 1.24E-05 |
| SerAGA.22 | -4.53803 | 1.05E-16 |
| SerAGA.37 | -4.60136 | 1.64E-16 |
| LeuTAG.5 | -4.61397 | 4.58E-07 |
| TyrGTA.43 | -4.71271 | 0.000363968 |
| SerGCT.9 | -4.78726 | 0.000569855 |
| ThrTGT.3 | -4.95799 | 3.88E-06 |
| CysGCA.9 | -4.9867 | 0.000443663 |
| GlnCTG.1 | -5.17113 | 0.000370386 |
| TyrGTA.63 | -5.55248 | 1.32E-07 |
| LeuTAG.10 | -5.74934 | 6.61E-12 |
| TyrGTA.75 | -5.77945 | 3.96E-07 |
| TyrGTA.69 | -5.96448 | 5.98E-05 |
| ProAGG.11 | -5.99826 | 5.71E-08 |
| TyrGTA.11 | -6.20296 | 3.56E-12 |
| TyrGTA.34 | -6.2482 | 1.65E-11 |
| TyrGTA.72 | -6.47593 | 1.56E-05 |
| GlnTTG.7 | -6.53276 | 2.01E-09 |
| TyrGTA.62 | -6.7874 | 3.37E-11 |
| GlnTTG.6 | -6.80242 | 1.52E-06 |
| TyrGTA.8 | -6.92138 | 6.10E-12 |
| TyrGTA.2 | -7.01583 | 3.56E-12 |
| TyrGTA.37 | -7.08318 | 1.73E-08 |
| ProTGG.33 | -7.08852 | 6.26E-10 |
| TyrGTA.7 | -7.13451 | 1.82E-13 |
| TyrGTA.39 | -7.23832 | 7.92E-14 |
| ProTGG.8 | -7.3484 | 3.41E-05 |
| TyrGTA.9 | -7.35098 | 1.60E-14 |
| TyrGTA.15 | -7.37364 | 7.65E-12 |
| TrpCCA.1 | -7.41409 | 1.78E-12 |
| TyrGTA.3 | -7.41545 | 5.16E-14 |
| TyrGTA.1 | -7.4546 | 2.94E-14 |
| TyrGTA.41 | -7.47747 | 6.62E-15 |
| TyrGTA.13 | -7.47883 | 3.45E-16 |
| TyrGTA.5 | -7.50459 | 2.75E-14 |
| TyrGTA.12 | -7.5431 | 1.11E-14 |
| TyrGTA.71 | -7.5461 | 2.53E-09 |
| TyrGTA.38 | -7.61067 | 2.53E-13 |
| TyrGTA.23 | -7.61644 | 2.34E-15 |
| TyrGTA.6 | -7.6245 | 4.01E-15 |
| TyrGTA.4 | -7.86068 | 9.02E-17 |
| TyrGTA.27 | -7.97605 | 9.80E-13 |
| TyrGTA.35 | -8.14863 | 1.07E-13 |
| TyrGTA.14 | -8.18338 | 3.60E-16 |
| TyrGTA.45 | -8.26412 | 4.81E-11 |
| TyrGTA.31 | -8.30471 | 3.64E-11 |
| TyrGTA.42 | -8.38152 | 1.23E-12 |
| GlnTTG.2 | -8.40242 | 1.67E-27 |
| TyrGTA.22 | -8.44592 | 5.52E-13 |
| SerCGA.4 | -8.54549 | 1.22E-16 |
| TyrGTA.49 | -8.54852 | 2.22E-13 |
| TyrGTA.10 | -8.5643 | 1.74E-16 |
| LeuCAG.2 | -8.6736 | 2.03E-13 |
| TyrGTA.53 | -8.73689 | 1.29E-15 |
| TyrGTA.16 | -8.82362 | 8.29E-19 |
| TyrGTA.18 | -8.86443 | 2.23E-11 |
| TyrGTA.19 | -8.93289 | 1.20E-15 |
| TyrGTA.64 | -9.02225 | 3.96E-17 |
| TyrGTA.55 | -9.08805 | 1.31E-15 |
| TyrGTA.44 | -9.18014 | 1.42E-16 |
| TyrGTA.51 | -9.18842 | 1.20E-16 |
| GlnTTG.1 | -9.20391 | 1.56E-25 |
| TyrGTA.29 | -9.24131 | 8.15E-18 |
| TyrGTA.56 | -9.31329 | 9.77E-20 |
| TyrGTA.52 | -9.33398 | 1.36E-18 |
| GlnTTG.8 | -9.38377 | 3.41E-28 |
| TyrGTA.46 | -9.43777 | 2.67E-14 |
| TyrGTA.20 | -9.67611 | 5.51E-18 |
| TyrGTA.33 | -9.70348 | 3.69E-13 |
| GlnTTG.3 | -9.82155 | 4.63E-26 |
| TyrGTA.50 | -10.1878 | 3.01E-18 |
| TyrGTA.24 | -10.2641 | 9.09E-23 |
| TyrGTA.48 | -11.1893 | 1.19E-17 |
| TyrGTA.47 | -11.398 | 3.24E-20 |
| TyrGTA.25 | -11.5015 | 1.11E-17 |
| GlnTTG.4 | -11.6281 | 6.33E-18 |
| TyrGTA.54 | -11.6416 | 2.08E-17 |
| TyrGTA.28 | -11.9044 | 1.10E-23 |
| TyrGTA.32 | -11.9692 | 4.61E-23 |
| ArgACG.7 | -12.1183 | 0.000969295 |
| ProAGG.10 | -17.1019 | 6.91E-37 |
