## Supplementary figures and images for "The small RNA component of Arabidopsis phloem sap and its response to iron deficiency"

### Supplemental Figure 1

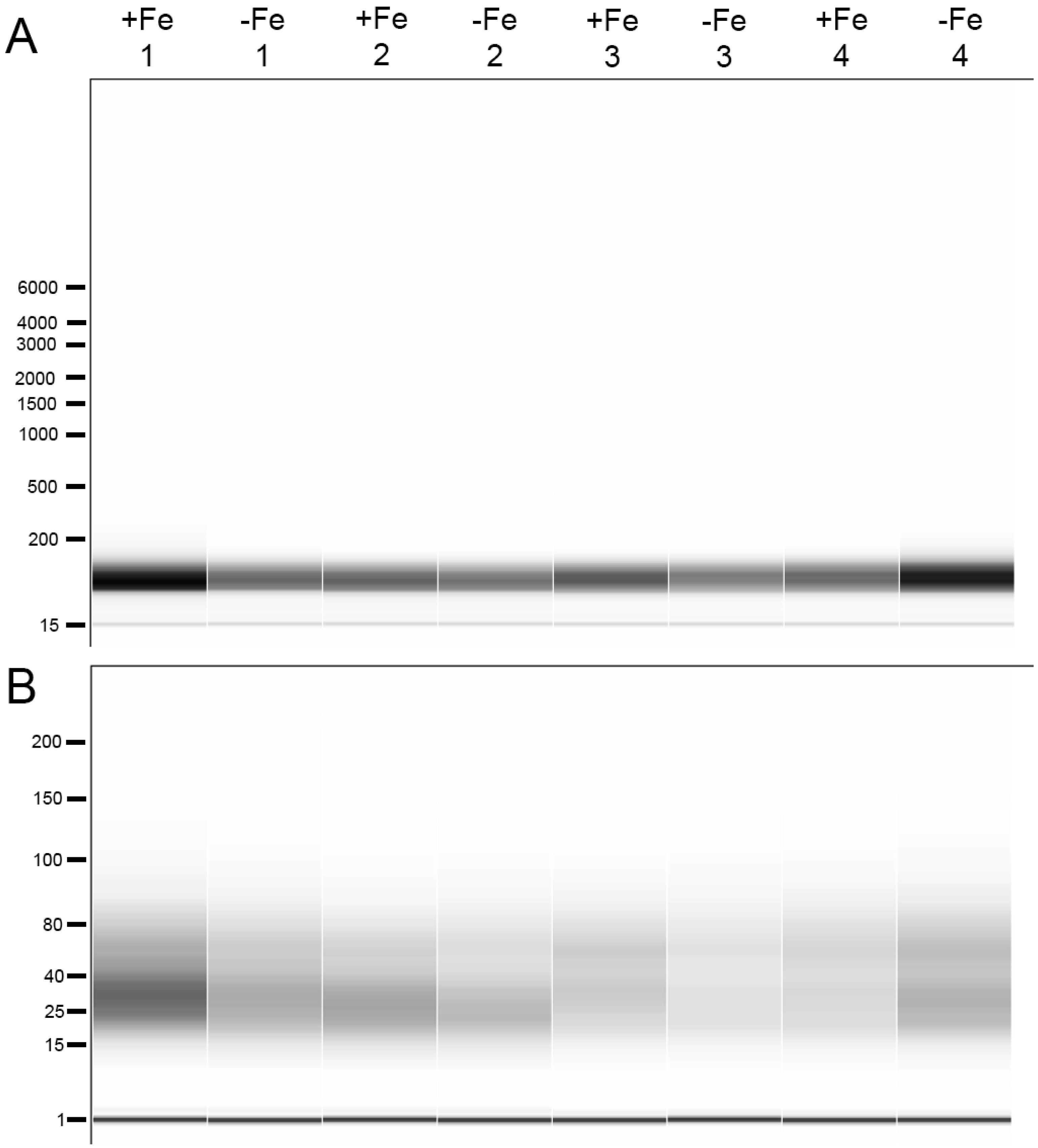

### Supplemental Figure 2

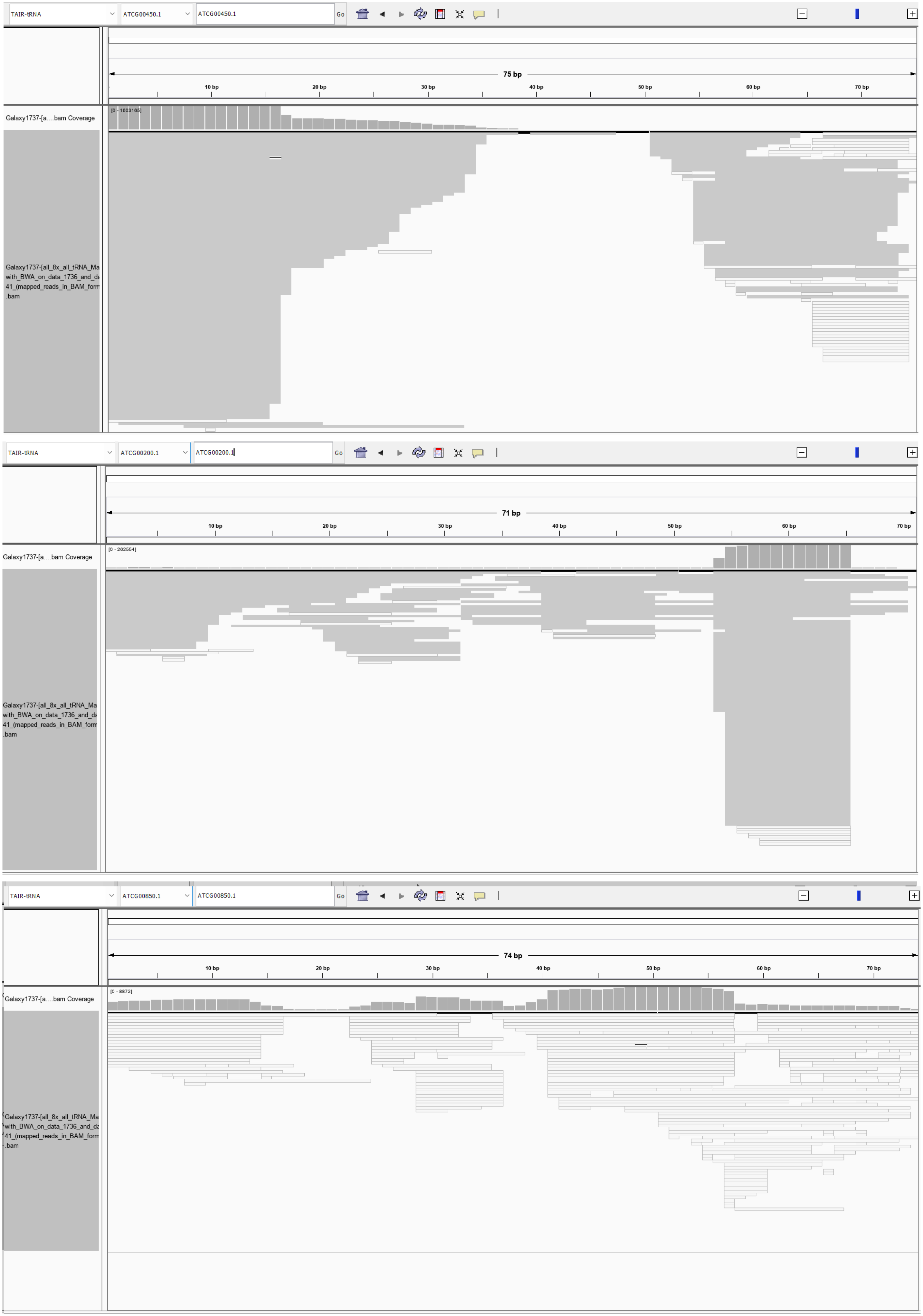

### Supplemental Figure 3

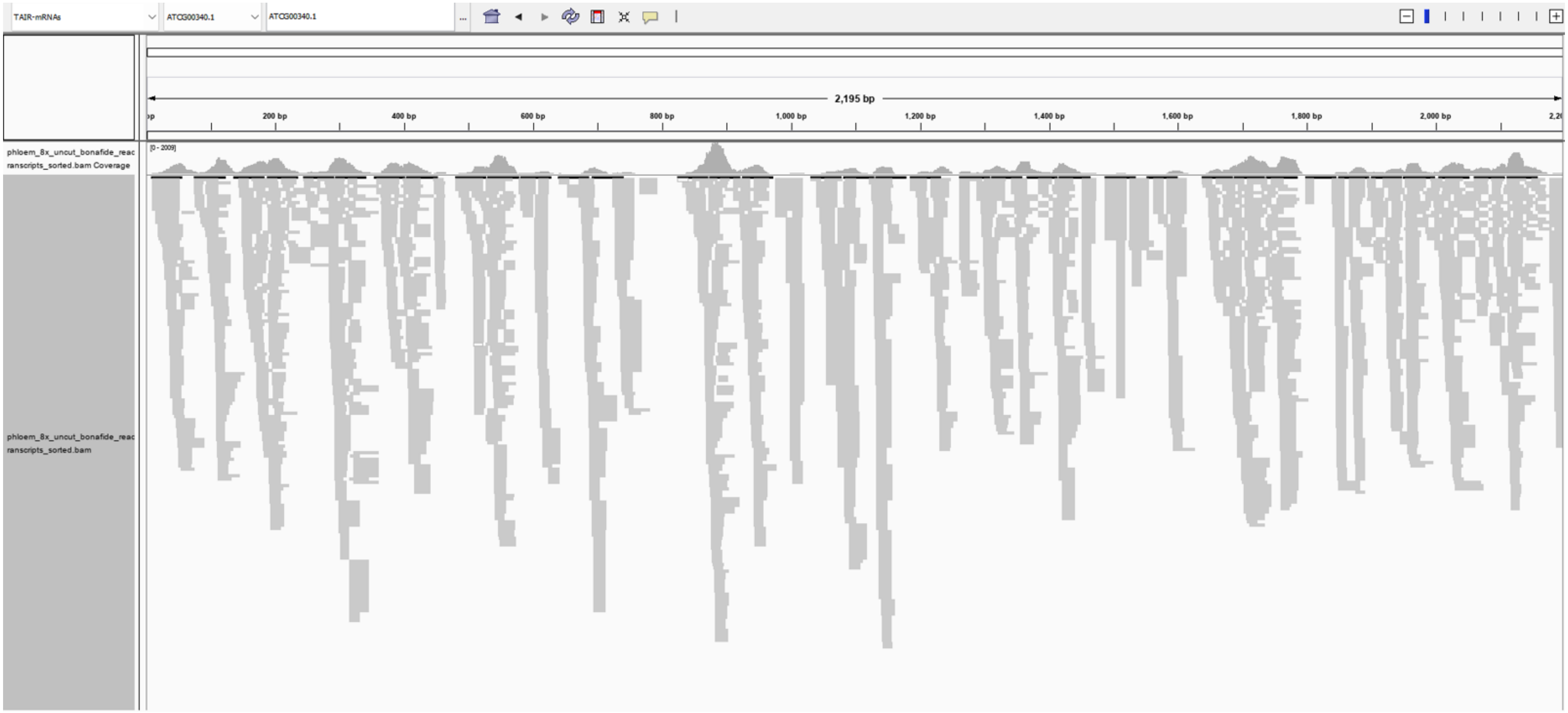

### Supplemental Figure 4

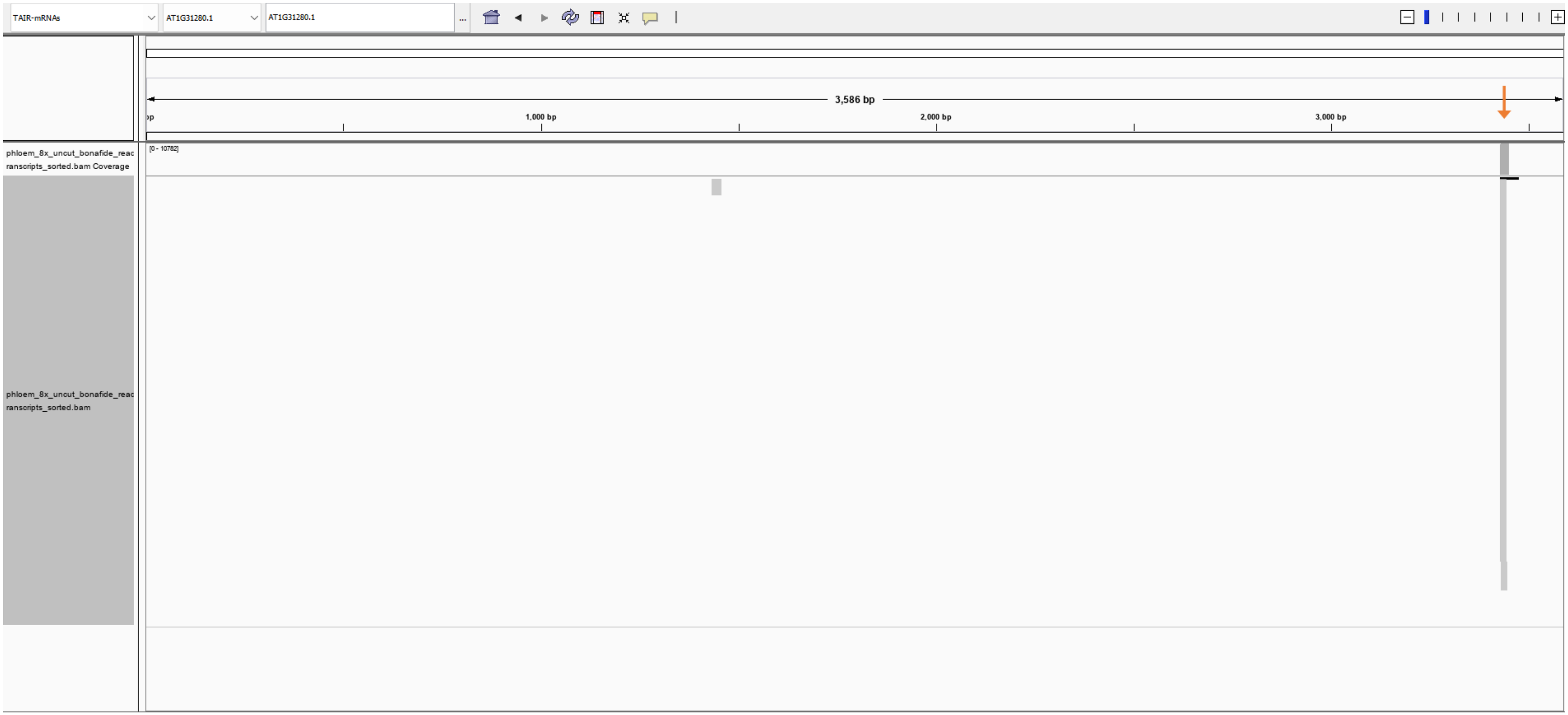
